# Validation of a non-appetitive effort-based foraging task as a measure of motivational state in male mice

**DOI:** 10.1101/2023.12.12.571234

**Authors:** Foteini Xeni, Caterina Marangoni, Megan G Jackson

## Abstract

Motivational disorders such as apathy syndrome are highly prevalent across neurological disorders but do not yet have an agreed treatment approach. The use of translational behavioural models can provide a route through which to meaningfully screen novel drug targets. Methods that utilise food deprivation in contrived environments may lack the sensitivity to detect deficits in self-initiated or self-driven behaviour. Animals monitored in more naturalistic environments may display more ethologically-relevant behaviours of greater translational value. Here, we aimed to validate a novel, non-appetitive effort-based foraging task as a measure of motivational state in mice. In this task, the mouse can freely choose to exert effort to forage bedding material and shuttle it back to a safe and enclosed environment. The amount of bedding material foraged is used as a readout of motivational state. Acute dopaminergic modulation with haloperidol, amphetamine and methylphenidate, and two phenotypic models known to induce motivational deficits (healthy ageing and chronic administration of corticosterone) were used to validate this task. Consistent with other effort-based decision-making tasks we find that foraging behaviour is sensitive to acute modulation of dopaminergic transmission. We find that both phenotypic models induce differing deficits in various aspects of foraging behaviour suggesting that the task may be used to parse different behavioural profiles from distinct disease phenotypes. Thus, without requiring extended training periods or physiological deprivation, this task may represent a refined and translational preclinical measure of motivation.

## Introduction

Developing our understanding of motivated behaviour is a critical aspect of behavioural neuroscience and pharmacology. Motivation encompasses a series of fundamental processes in which an organism interacts with its environment in pursuit of a goal. Disruption to these processes can have a profound clinical impact. For example, apathy syndrome is a psychiatric syndrome characterised by a reduction in self-initiated behaviour and emotional blunting (Marin, 1991, Levy and Dubois, 2006). It is pervasive across numerous neurological and neurodegenerative diseases, yet does not currently have an agreed treatment approach (Benoit et al., 2008, den Brok et al., 2015). This may be in part be due to a lack of understanding of its underlying neurobiology, and a preclinical pipeline through which to screen novel drug targets.

Our currently available preclinical methods for assessing goal-directed behaviour derive from operant conditioning paradigms such as the effort for reward task (EfR) or progressive ratio task (Salamone et al., 1994, Richardson and Roberts, 1996). Here, the rodent is food restricted, and learns to make an effortful number of operant responses (lever press or nose poke) to receive a palatable food reward. In the case of the EfR task, an extra dimension of choice is added, where the rodent can choose to exert high effort for a higher value reward, or consume the lower value freely available lab chow. The EfR task has been valuable in understanding the neural substrates underlying reinforcement-based goal-directed behaviour, demonstrating a key role for the mesolimbic dopaminergic (DA) system alongside evidence for other interacting systems across species (Salamone et al., 1994, Nunes et al., 2013, Salamone et al., 2018, Marangoni et al., 2023). However, we must be mindful that such operant tasks take weeks-months to train, are run in a contrived environment, with sometimes high levels of food restriction. The purpose of this is to induce a robust and well-defined behavioural response, but may come at a cost to behavioural interpretation in some cases. For example, it is unclear whether a deficit in intrinsic self-drive characteristic of apathy can be detected under the pressure of a significant external motivator such as hunger induced by food restriction. Food restriction has also been shown to fundamentally change physiology and brain function, which may impinge on our neural circuits of interest in an unknown way (Bubenik et al., 1992). There is a growing push towards the use of ethological, evolutionarily-conserved behaviours as a focal point of behavioural tasks, and may be of particular value where self-driven behaviours are the focus of the study (Puścian and Knapska, 2022). By utilising spontaneous, intrinsically rewarding behaviours, we can potentially obtain robust, reliable responses in more complex environments without the need to externally modulate motivational state via food restriction. In this way, motivational state can be assessed in a potentially more translatable context. In addition, by removing the need to physiologically deprive the rodent and reducing training time, these tasks provide the opportunity to refine rodent lifetime experience in the lab and thus improve welfare.

In this work, we present a novel, non-appetitive behavioural task which draws on the mouse’s intrinsic drive to forage for bedding material. The apparatus is designed to encourage the mouse to exert effort to forage bedding material and shuttle it back to a safe and enclosed environment. The amount of bedding material foraged is used as a readout of motivational state. This is distinct from nest building assays where the quality of nest is scored over prolonged periods of time. Here we capture the activational foraging aspect of this behaviour in time windows conducive to drug screening. Through acute modulation of the dopaminergic system, and the use of phenotypic models previously shown to induce a deficit in reward motivation (healthy ageing and chronic corticosterone (CORT) administration (Dieterich et al., 2019, Jackson et al., 2021)) we aimed to validate the effort-based foraging (EBF) task as a refined measure of motivational state in mice. The D2 receptor antagonist haloperidol was selected to reduce DA transmission and has been used to validate previous effort based decision making studies (Salamone et al., 1994, Marangoni et al., 2023). The psychostimulant amphetamine and dopamine transporter inhibitor methylphenidate were similarly selected to elevate DA transmission and have also been utilised in the validation of motivation-based tasks (Marangoni et al., 2023).

### Methods Subjects

7 cohorts of male C57BL/6 mice (Envigo (UK)) were used in these experiments (**see S1 for a summary of ages, n numbers and cohorts used per experiment**) and were all housed, handled and habituated under the same conditions. Mice were singly housed in enriched open-top Techniplast 1284 conventional cages to avoid significant levels of in-fighting commonly observed with this strain and sex and thus minimise any impacts of aggression and associated psychosocial stress. Mice underwent twice daily health checks to ensure good welfare throughout experimental testing. Each cage was enriched with a small red house, a wooden chew, a tube, and a tube hung from the ceiling. Standard laboratory chow (Purina, UK) and water were provided *ad libitum* throughout experiments. Mice were housed in a 12:12 reverse light schedule, where lights turn OFF at 8.15 am, and turn ON at 8.15 pm. Testing did not begin until all mice showed no aversive behaviours to cup handling (∼ 3 days). All behavioural testing took place in the animal’s active phase (ZT (Zeit time) 13-21 (9.15 am-17.15 pm)). Sample sizes were based on detecting a large effect size with mean difference and variance based on previous behavioural studies using a similar task and acute pharmacological manipulations and phenotypic models (Marangoni et al., 2023, Jackson et al., 2021). In all pharmacological studies the experimenter was blind to treatment. It was not possible to be blinded during the healthy ageing study due to clear phenotypic differences between groups. In the case of acute pharmacological studies and studies relating to changes in the external environment mice were randomly assigned a treatment group using a within-subject counterbalanced Latin square design. The experimental unit for all studies was the individual animal. All experiments were performed in accordance with Animals Scientific Procedures Act (ASPA, UK) 1986 and were approved by the University of Bristol Animal Welfare and Ethical Review Body (AWERB). All equipment developed and used in this work including materials, 3D print files and supporting code will be made available upon acceptance. Arena dimensions are provided in **S2**.

### Effort-based foraging task arena

A foraging arena was developed in house by MGJ to assess non-appetitive forage based motivation in mice. The apparatus is designed to encourage the mouse to exert effort to forage bedding material and shuttle it back to a safe and enclosed environment. The standard arena set up is outlined in **fig 1**. Amended versions of the task for affective reactivity testing and effort curve testing (see further details below) are outlined in **fig 2**.

**Figure 1.**
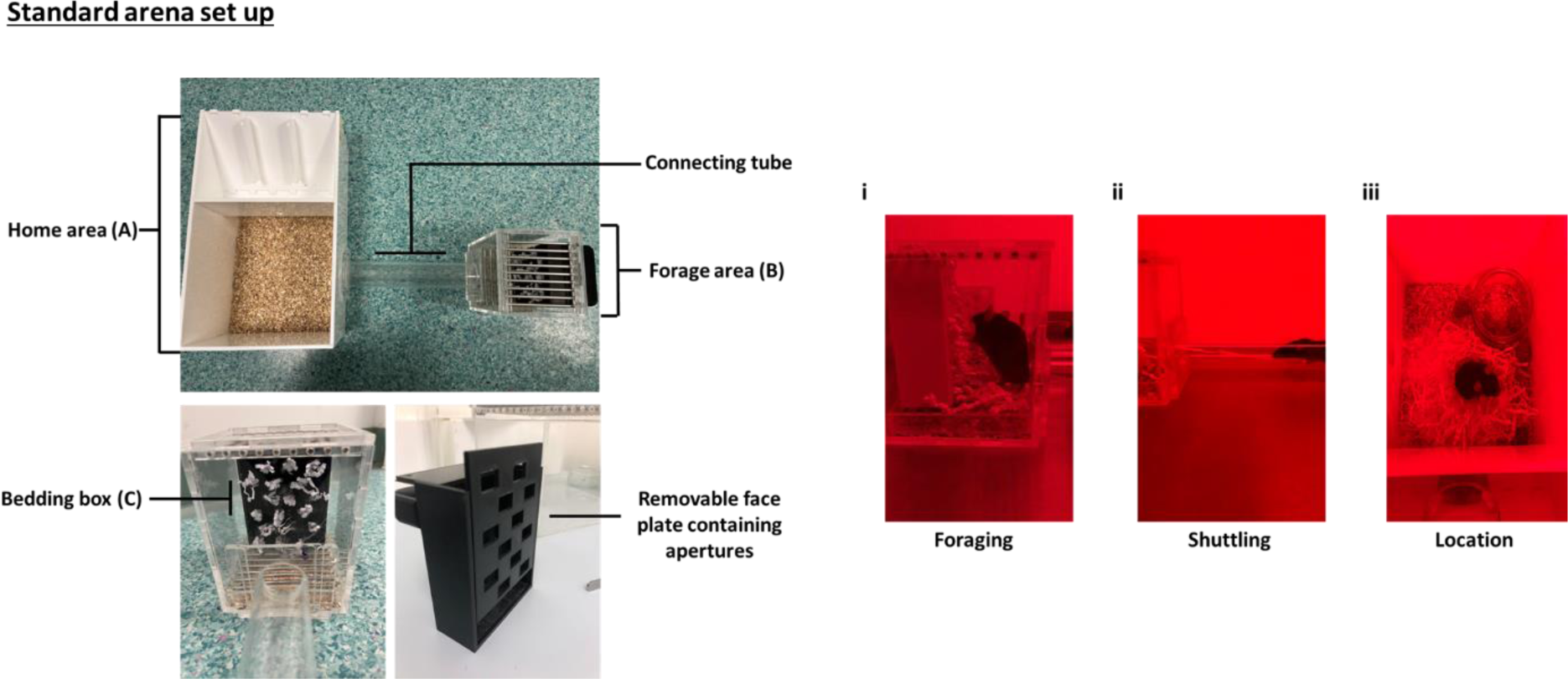
Standard effort-based forage arena set up. The arena consists of four main components. The home area (**A**), the connecting tube, the forage area (**B**) and the bedding box (**C**). The home area contains woodchip on a flat surface, and free access to food and water. It is covered by a detachable lid when in use. The use of solid Perspex creates a safe, enclosed environment. The forage area contains a barred floor, with woodchip underneath. It is covered with a detachable barred lid while in use. The bedding box is fixed to the back of the forage area with magnet bar, which is external to the arena. The bedding box is filled with 18 g of Sizzlenest. Throughout the task period, the mouse has the option to traverse the tube, forage bedding from the bedding box by pulling it with their mouth/paws (i), and shuttle it back through the tube (ii) to the home area (iii). The use of a barred floor in the forage area in combination with the more ‘open’ environment (use barred lid to allow air flow and clear Perspex) promotes shuttling of bedding to the home area. In some cases, the mouse may forage bedding (successfully pull bedding from the box) but leave some in the forage area. The main outputs from the task are the amount of bedding pulled from the bedding box and shuttled to the home area, and the *% of foraged bedding that is taken to the home area*.

**Figure 2.**
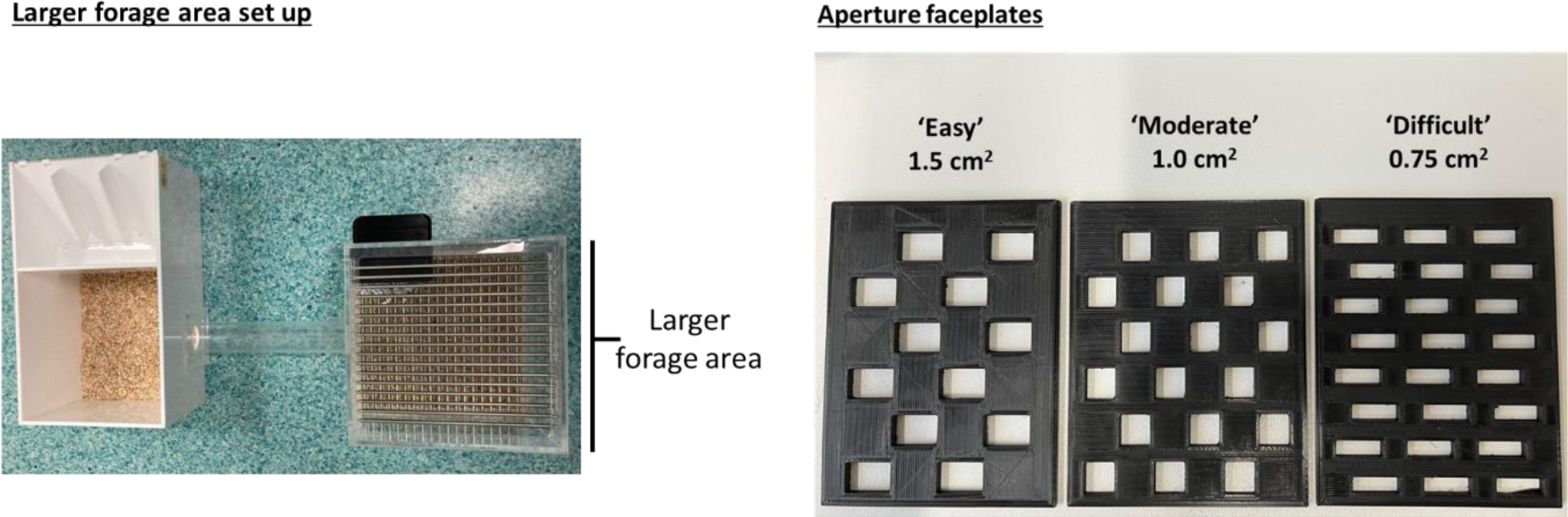
Amended arena set ups. In an amended arena set up, a larger forage area is used to assess the impact of stress induced by aversion to open spaces on foraging behaviour (affective reactivity). The bedding box is placed nearer the tube entrance to ensure that no additional shuttling distance induced by placing the bedding box at the back impacts on behaviour. To modulate effort required to forage bedding during ‘effort curve’ testing, three different face plates containing different size apertures are used. The moderate aperture is used consistently for acute pharmacological testing.

### Habituation to the effort-based foraging task

Mice were placed in the home compartment of the forage arena (**fig. 1**, part A) containing woodchip only and were left to explore for a period of 10 minutes. On the second and third day, they were left for a period of 5 minutes. Woodchip was changed between mice. On the fourth day, they were placed in the forage arena for 4 hours. This time, the arena contained a bowl of standard lab chow and their water bottle placed in the main compartment, and a bedding box placed in the foraging area (**fig 1**, part B). The bedding box was filled with 18g of white sizzlenest (Datesand), which could be pulled through 1.5cm^2^ apertures. Amount of bedding (g) pulled out of the bedding box and shuttled to the main compartment was measured. Bedding pulled out but left in the forage compartment was also measured. Across all studies the main output measures were total bedding shuttled to the home area (g) and percentage bedding foraged taken to the home area. Terms relating to behavioural output are outlined in **table 1**.

**Table 1.** Description of key behavioural output terms.

| Term | Description |
| --- | --- |
| **Total bedding shuttled to the home area (g)** | The amount of bedding pulled (foraged) from the bedding box and taken through the tube to the home area. |
| **Percentage bedding foraged taken to the home area (%)** | Of the total bedding pulled (foraged) from the bedding box, how much is transported to the home area instead of left on the floor of the forage area, expressed as a percentage. |

### Acute pharmacology

N = 16 mice aged 12 −14 weeks were used for the haloperidol and i.p amphetamine experiments, and n = 16 mice aged 13 - 15 weeks were used for the methylphenidate and oral amphetamine studies. Following habituation mice underwent three 2 hour sessions of foraging using the bedding box face plate with 1 cm^2^ apertures which produced a moderate level of difficulty. Sessions were repeated to ensure stability in performance before starting drug testing.

An acute study consisted of four 2-hour test sessions (vehicle and 3 doses). Each test session was separated by at least 2 days to ensure drug washout between sessions (**fig 3**). Each mouse received each dose in a within-subject, counterbalanced design. Drugs were administered in a 10 ml/kg dose volume. Haloperidol (0.01 – 0.1 mg/kg, i.p, Sigma UK), amphetamine (0.1 – 0.3 mg/kg, oral and i.p, Sigma UK and methylphenidate (1, 3 and 10 mg/kg, oral, Sigma UK) were used in this study. Drugs, doses, route of administration, pre-treatment time and vehicles are outlined in **table 2**. Doses and pre-treatment times were selected based on available pharmacokinetic data and previously used doses (Marangoni et al., 2023, Stuart et al., 2017, Griesius et al., 2020). Mice were restrained and scruffed for i.p injection using a refined protocol outlined in (Davies et al., 2022). In the case of oral dosing, mice were trained to drink voluntarily from a syringe. Briefly, mice were first habituated to a 20 % condensed milk solution (Nestlé Carnation) by putting ∼ 50 µl on a surface in the home cage and letting them voluntarily consume it in their own time. The next day mice were given 300 µl condensed milk solution via a 1 ml syringe through the bars of the lid of the home cage. This was repeated until the mouse would approach the syringe and consume the solution in ∼< 1 minute. On average this took 3 days. Using this method, all mice consumed all drug in a stress-free manner. Drugs were dissolved in water, and administered via a syringe in a final solution of 20% condensed milk.

**Figure 3.**
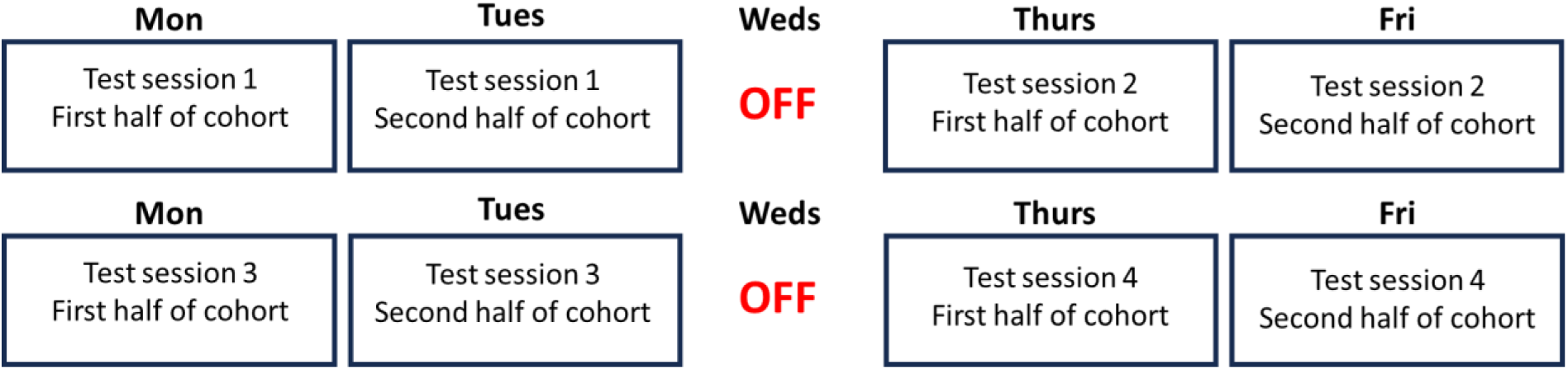
Test session design. In a 3 dose plus vehicle design, testing takes place over 4 testing sessions (two weeks). Each test session is separated by at least 2 days to ensure drug washout between sessions.

**Table 3.**
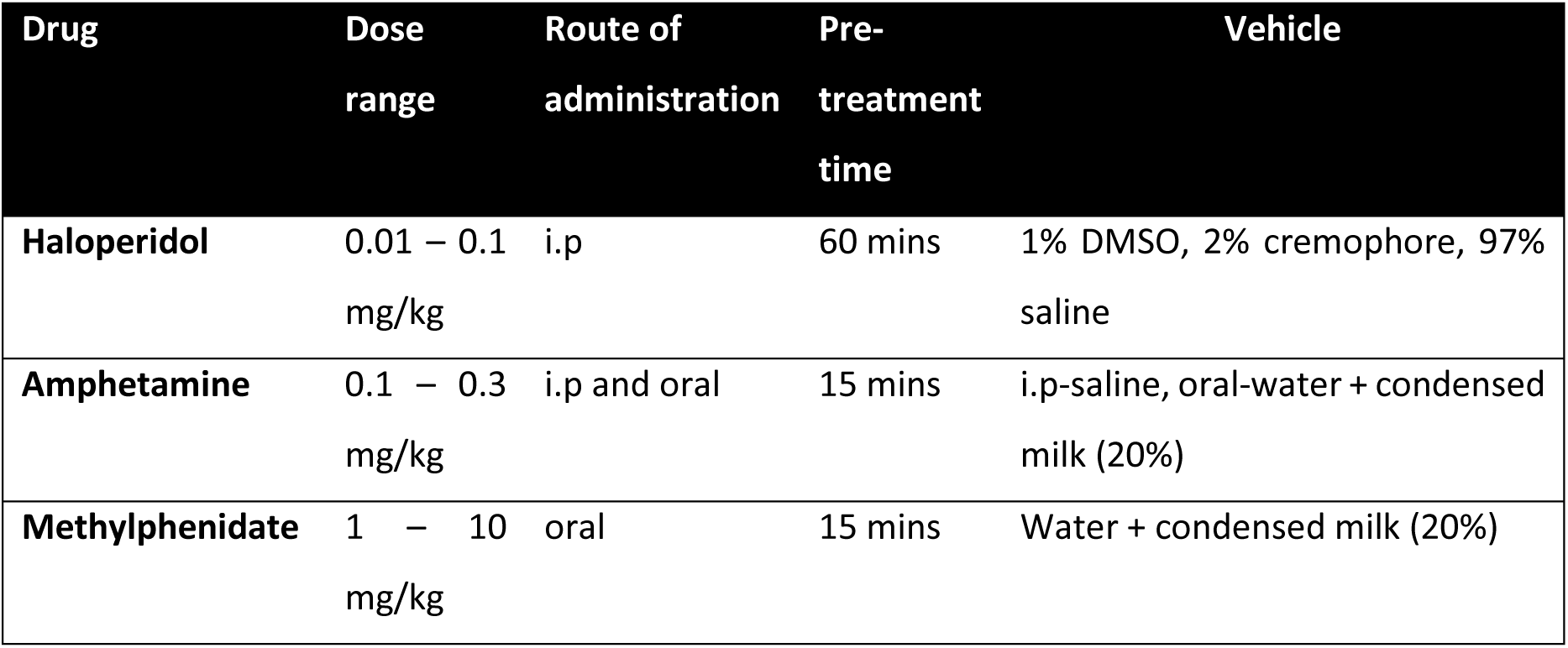
Summary of acute drug studies. i.p – intraperitoneal, DMSO- dimethyl sulfoxide. Oral indicates voluntary ingestion of substance via syringe.

### Activity sensing

To check for changes in general locomotor activity following treatment with haloperidol and amphetamine, activity was monitored in the foraging area using a passive infrared sensor system. The system was built in-house by MGJ to monitor home cage activity in individually-housed mice, based on a system previously developed by (Brown, 2016). 4 activity sensors with built-in amplifiers (AMN 2,3,4 series Motion Sensor, Panasonic) were read using an Arduino Mega 2560 with an Arduino Mega daughter board (SchmartBoard, Mouser). Data was read onto an SD card as CSV file using an SD card adaptor (TFT LCD w/microSD Breakout, Adafruit) and exact time was outputted alongside the data using a RealTime clock (Clock & Timer Development Tools PCF8523 RTC for RPi, Adafruit). Each sensor was placed above the middle of the forage area, approximately 5 cm from the top of the cage. Movement was detected by the sensor every 100 msec. A percentage of movement within 10 secs was calculated and outputted in 10 second time bins.

### Phenotypic models

#### Chronic corticosterone

In two cohorts of 16 mice run consecutively, half were allocated to the treatment group (corticosterone, HBC complex (2.5mg/100ml), Sigma Aldrich, UK) in drinking water and half to the non-treatment group (standard drinking water). Mice on average drank ∼ 5 ml water/day and weighed on average 35 g. Mice were treated for 3 weeks before undergoing the EBF task. Following standard habituation including an initial foraging session, mice underwent three 2 hour foraging sessions under different effort contingencies governed by the size of the bedding box aperture, allowing the formation of an ‘effort curve’. The aperture size was changed by swapping the face plate on the bedding box. The aperture sizes were 1.5 cm^2^ (easy), 1 cm^2^ (moderate) and 0.75 cm^2^ (difficult). While the individual aperture differed in size, the total available surface area for foraging remained the same between face plates by increasing the number of apertures available (**fig 2**). Each mouse underwent foraging under each condition in a within-subject, counterbalanced design.

Mice also underwent two x 2 hour foraging sessions using the 1.5 cm^2^ aperture where the bedding box is placed in either the enlarged or standard forage area (**fig 2**). This was to test affective reactivity driven by mice finding more open, novel spaces aversive. Mice underwent both sessions at least two days apart in a within-subject counterbalanced design.

#### Healthy ageing

Following standard habituation, n = 12 young (13 - 23 weeks old) and n = 11 aged mice (44 - 54 weeks old) underwent the effort curve and affective reactivity testing as described above. To account for potential age-related changes in motoric ability confounding aged mouse performance, the task was extended to 4 hours in this case.

### Environmental changes

#### Free bedding

In a counterbalanced, within-subject design, n = 16 mice (35 weeks old) underwent two 2 hour forage sessions using the 1 cm^2^ (mid-size) aperture with either the presence or absence of 10 g of freely available sizzle bedding, placed in the main compartment where bedding is typically transported across the course of the session. Total bedding foraged from the bedding box and amount left in the forage compartment was analysed.

#### Warming

A heat mat (NEKOSUKI reptile heating mat) was placed under the main compartment of the forage arena, and the floor was warmed to a temperature of ∼29 °C, which has been shown to be the threshold for the thermoneutral zone for mice (Vialard and Olivier, 2020). n = 12 mice (30 weeks old) were habituated to the warmed arena in a 20 minute session without the bedding box the day preceding the test sessions. In a counterbalanced, within-subject design, the mice underwent two 4 hour test sessions (+/- warming).

### Statistical analysis

Data were tested for normality using a Shapiro-Wilk test. Where data were normally distributed and required single factor analysis, a repeated measure (RM) one-way ANOVA (Geisser-Greenhouse corrected) was used. Where significant main effects were observed (p < 0.05), Dunnett’s post-hoc comparisons were carried out. Where data violated normality, a Friedman test was applied. Where a significant main effect was observed, Dunn’s post hoc comparisons were carried out. Trend level main effects (p < 0.1) are reported but not further analysed. Where two factor analysis was required (age*aperture, age*arena size, CORT treatment*aperture, CORT*arena size), a RM two-way ANOVA was applied. Where significant main effects occurred, these were tested using Sidak post-hoc comparisons. Outliers were defined as data points ± 2SDs away from the mean. Outliers were replaced with the group mean to permit repeated measures analysis. If data was missing due to random errors in recording then a mixed model was applied instead. All instances of outlying/missing data points are provided in **S3**. Data were graphed and analysed using GraphPad Prism v 10.0.03.

## Results

### Foraging behaviour is sensitive to acute dopaminergic modulation

Acute administration of haloperidol (0.01 – 0.1 mg/kg) had a main effect on total bedding shuttled (X2 = 17.15, p = 0.0007, Friedman Test). Dunn’s post-hoc test revealed 0.1 mg/kg reduced bedding shuttled (p = 0.0005) (**fig. 4A**). To assess a potential interaction between drug effect and intrinsic motivational state, the cohort was then stratified into ‘high’ and ‘low’ performers using a median split of bedding foraged under vehicle conditions. There was a main effect of drug (F(2.307, 29.99) = 11.84, p < 0.0001 RM two-way ANOVA), main effect of motivational state (F(1,13) = 32.73, p < 0.0001) and a drug*motivational state interaction (F(3,39) = 4.641, p < 0.0001). Sidak’s post hoc analysis showed that in both low and high motivational groups, 0.1 mg/kg reduced bedding shuttled (p = 0.0051 and p = 0.0065 respectively) (**fig. 4B**). Analysis of % of foraged bedding taken to the home area revealed no effect of drug (p > 0.05) (**fig. 4C**). There was also no effect of drug on general locomotor activity in the forage area (p > 0.05) (**fig. 4D**).

**Figure 4.**
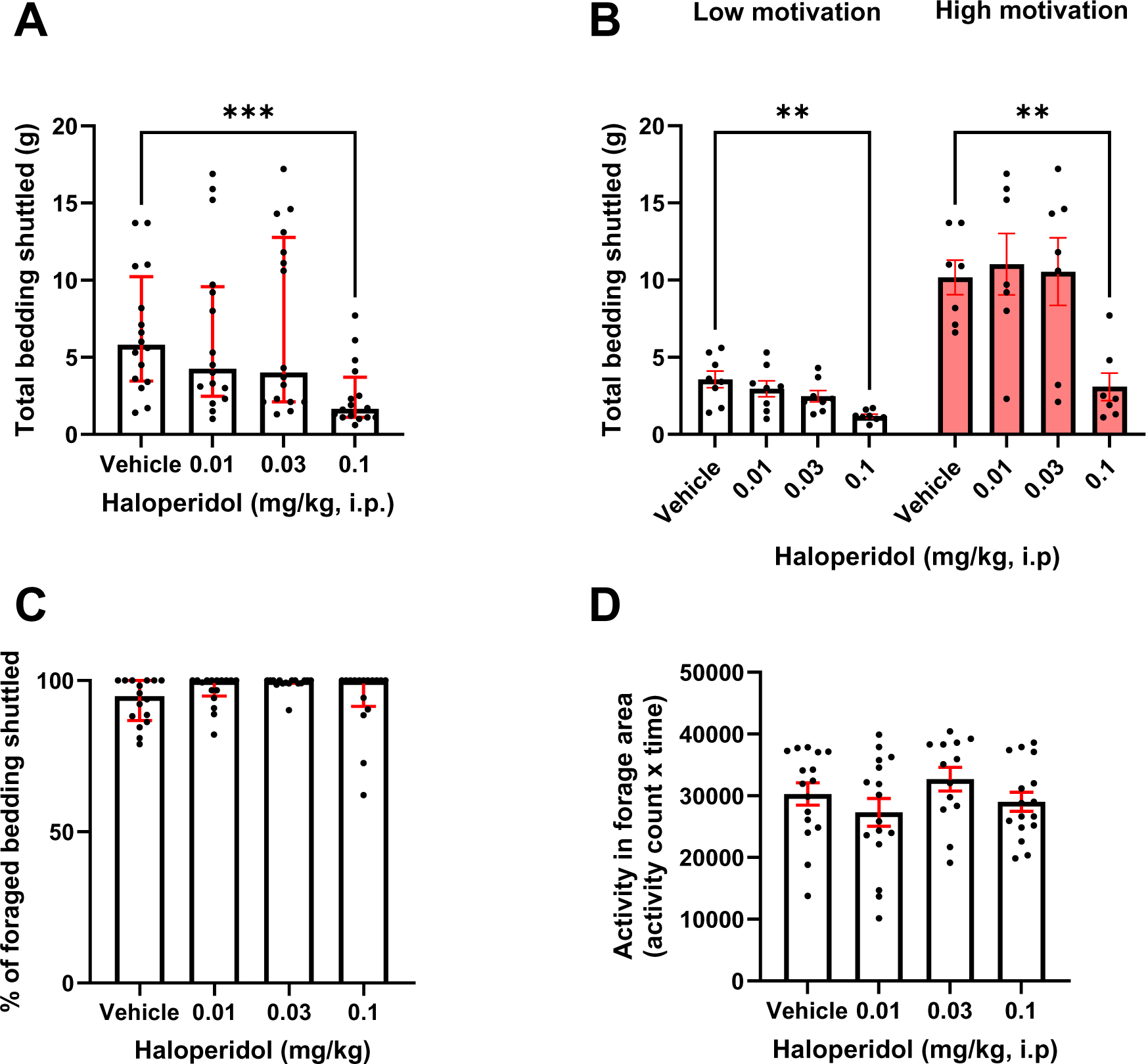
Haloperidol reduces foraging behaviour without changing general activity. Mice were administered haloperidol and put in the EBF task for 2 hr. **A** 0.1 mg/kg reduced total bedding shuttled (p < 0.001). **B** This effect was consistent across low and high performers (p < 0.01). **C** There was no effect of haloperidol on % of foraged bedding taken to home area and **D** general locomotor activity in the forage area. Bars are mean ± SEM or median ± interquartile range where data are non-parametric (A, C). **p< 0.01, ***p<0.001.

Acute administration of methylphenidate (1 – 10 mg/kg) also had a main effect on total bedding shuttled (F(23.62,35.42) = 7.353, p = 0.0013, RM one-way ANOVA). Dunnett’s post hoc analysis showed that 10 mg/kg increased bedding shuttled (p = 0.0003) (**fig. 5A**). Mice were then stratified by intrinsic motivational state as described above to assess drug and motivational state interaction. There was a main effect of drug (F(2.398,36.37) = 10.15, p = 0.0001, a main effect of motivation (F(1,14) = 26.61, p = 0.0001) and a drug*motivational state interaction (F(3,42) = 4.992, p = 0.0047) on bedding shuttled. Sidak’s post hoc analysis showed that in the lower motivational state group, both 1 and 10 mg/kg increased bedding shuttled (p = 0.0272 and p = 0.0044 respectively), while in the high group only 10 mg/kg increased foraging behaviour (p = 0.0161) though there was a trend towards 3 mg/kg additionally increasing foraging (p = 0.0581) (**fig. 5B**). There was no effect of drug on % of foraged bedding brought to home area (p > 0.05) (**fig. 5C**).

**Figure 5.**
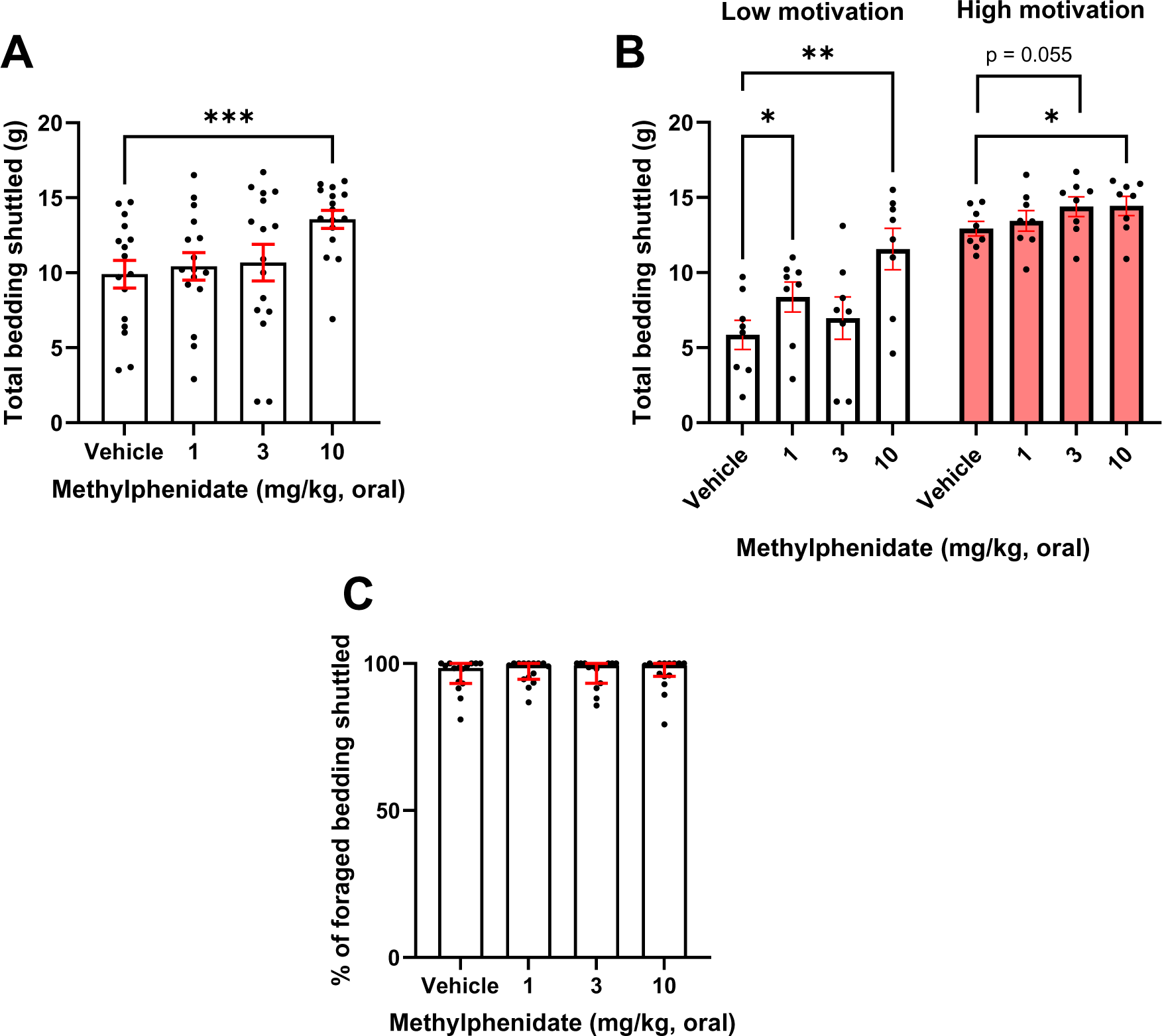
Methylphenidate increases foraging behaviour. Mice were administered methylphenidate and put in the EBF task for 2 hr. **A** 10 mg/kg increased total bedding shuttled (p < 0.001). **B** In low performers, both 1 and 10 mg/kg increased bedding shuttled (p < 0.05 and p < 0.01 respectively) but in high performers just 10 mg/kg increased bedding shuttled (p < 0.05). **C** There was no effect of methylphenidate on % of foraged bedding taken to home area. Bars are mean ± SEM or median ± interquartile range where data are non-parametric (C). *P< 0.05, **p< 0.01, ***p<0.001.

Amphetamine was administered both orally and via i.p in separate experiments (0.1 – 1 mg/kg). When administered i.p, there was a main effect of drug on bedding shuttled (F(2.836, 42.54) = 3.541, p = 0.0243). Dunnett’s post hoc analysis showed that 0.3 mg/kg decreased bedding shuttled (p = 0.0122) and a trend towards 0.1 mg/kg exerting a similar effect (p = 0.0607) (**fig. 6A**). When stratified by motivational state, there was a main effect of drug on bedding shuttled (F(2.061, 28.86) = 4.767, p = 0.0155). There was also a main effect of motivational state (F(1,14) = 13.46, p = 0.0025) and a drug*motivational state interaction (F(3,42) = 3.065, p = 0.0382). Sidak’s post hoc analysis revealed that these drug effects were specific to the high performance group, where both 0.1 mg/kg and 0.3 mg/kg decreased bedding shuttled (p = 0.0443 and p = 0.0049 respectively) (**fig. 6B**). There was no effect of drug on % of foraged bedding taken to the home area (p > 0.05) and no general effect of the drug on locomotion in the foraging area (p > 0.05) (**fig. 6C&D**). In contrast, amphetamine administered orally had no effect on bedding shuttled (p > 0.05) (**fig. 7A**) and similarly no interaction when stratified by motivational state (p > 0.05), though there was a main effect of motivational state as expected (F(1,14) = 4.813, p = 0.0456) (**fig. 7B**). Additionally there was no effect of drug on % of foraged bedding taken back to the home area (p > 0.05) (**fig. 7C**).

**Figure 6.**
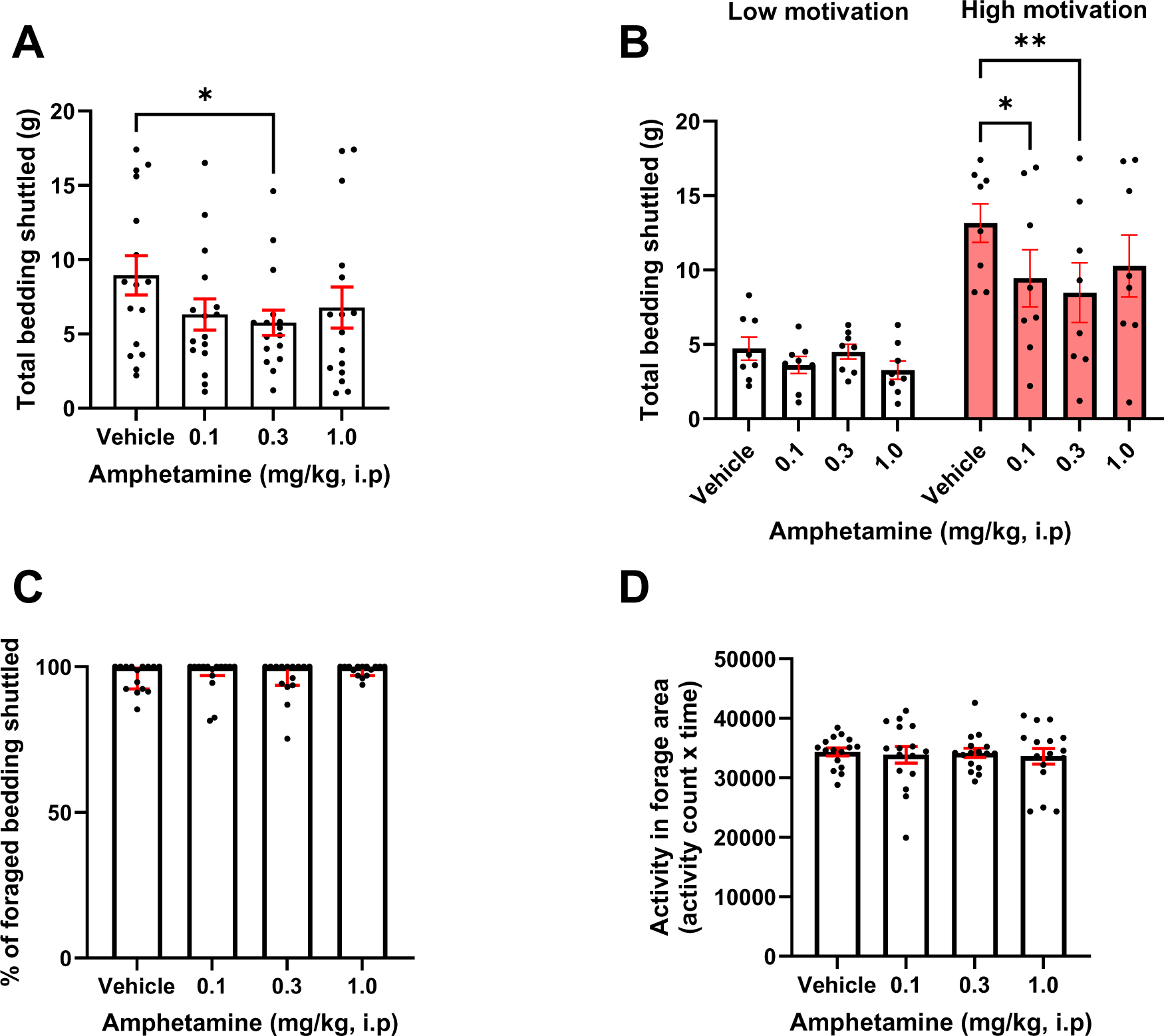
i.p administration of amphetamine reduces foraging behaviour without changing general activity. Mice were administered amphetamine and put in the EBF task for 2 hr. **A** 0.3 mg/kg reduced total bedding shuttled (p < 0.05). **B** This effect was specific to high performers, where both 0.1 and 0.3 mg/kg reduced bedding shuttled (p < 0.05 and p < 0.01 respectively). **C** There was no effect of amphetamine on % of foraged bedding taken to home area and **D** general locomotor activity in the forage area. Bars are mean ± SEM or median ± interquartile range where data are non-parametric (C). *p< 0.05, **p< 0.01, ***p<0.001.

**Figure 7.**
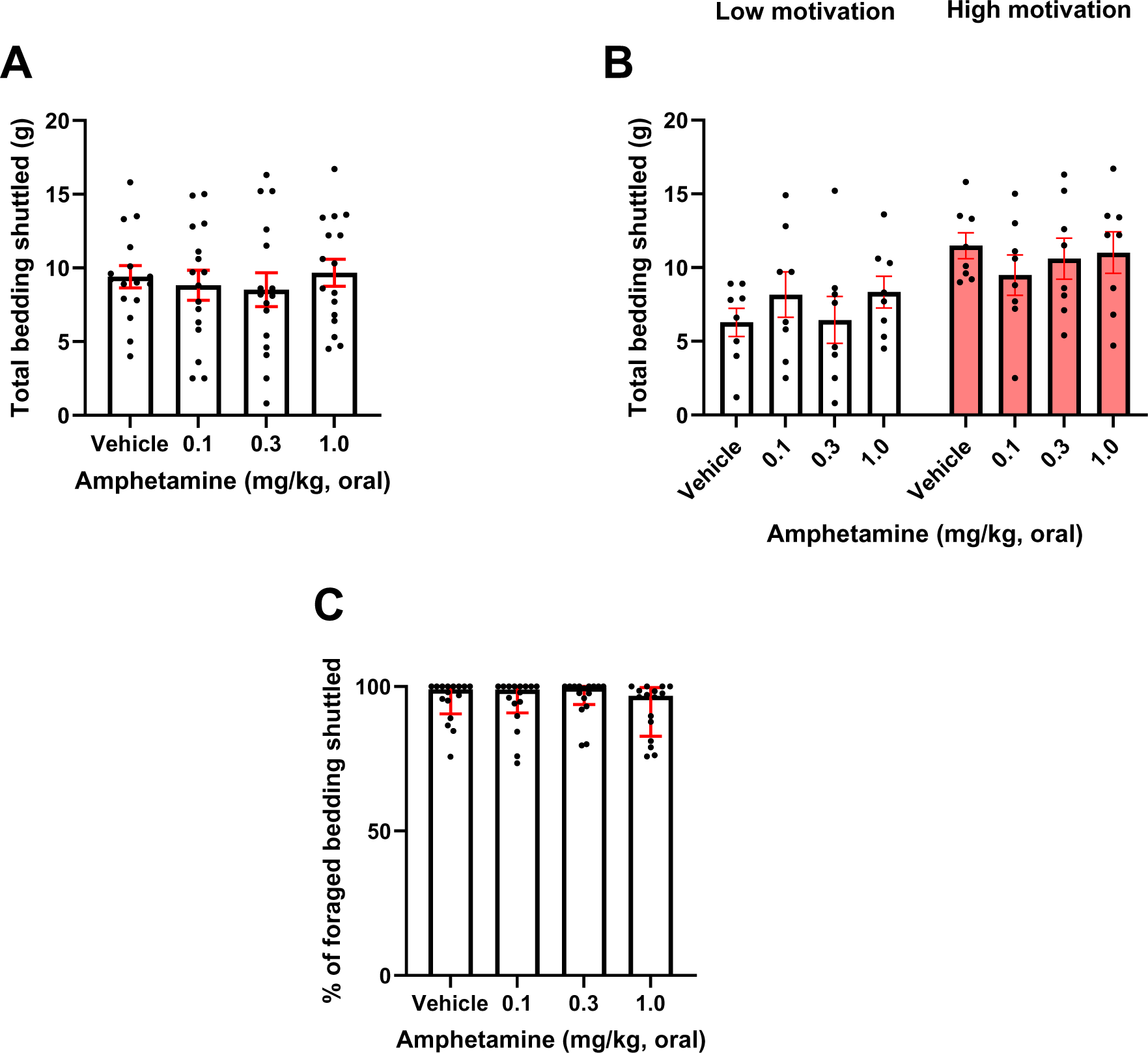
Oral administration of amphetamine has no effect on foraging behaviour. Mice were administered amphetamine orally and put in the EBF task for 2 hr. **A** There was no effect of drug on total bedding shuttled (p > 0.05). **B** There was no interaction between drug effect and motivation population. **C** There was no effect of haloperidol on % of foraged bedding taken to home area. Bars are mean ± SEM or median ± interquartile range where data are non-parametric (C).

### Foraging behaviour is sensitive to phenotypic models of motivational deficit

Young and aged mice underwent a series of experiments in the EBF task. Exploration of the forage area during habituation was compared between age groups across sessions to ensure potential deficits in further experiments were not driven by this factor, which has previously been reported (Jackson et al., 2021). There was a main effect of age (F(1,21) = 10.63, p = 0.0037), a main effect of session (F(1.904, 39.98) = 27.93, p < 0.0001) and an age*session interaction (F(2,42) = 3.656, p = 0.034). Sidak’s post hoc analysis revealed that young mice show greater levels of exploration of the forage area in the first two habituation sessions (p = 0.0053 and p = 0.0083) respectively, however this gap closes by the third session (p > 0.05) (**fig. 8A**).

**Figure 8.**
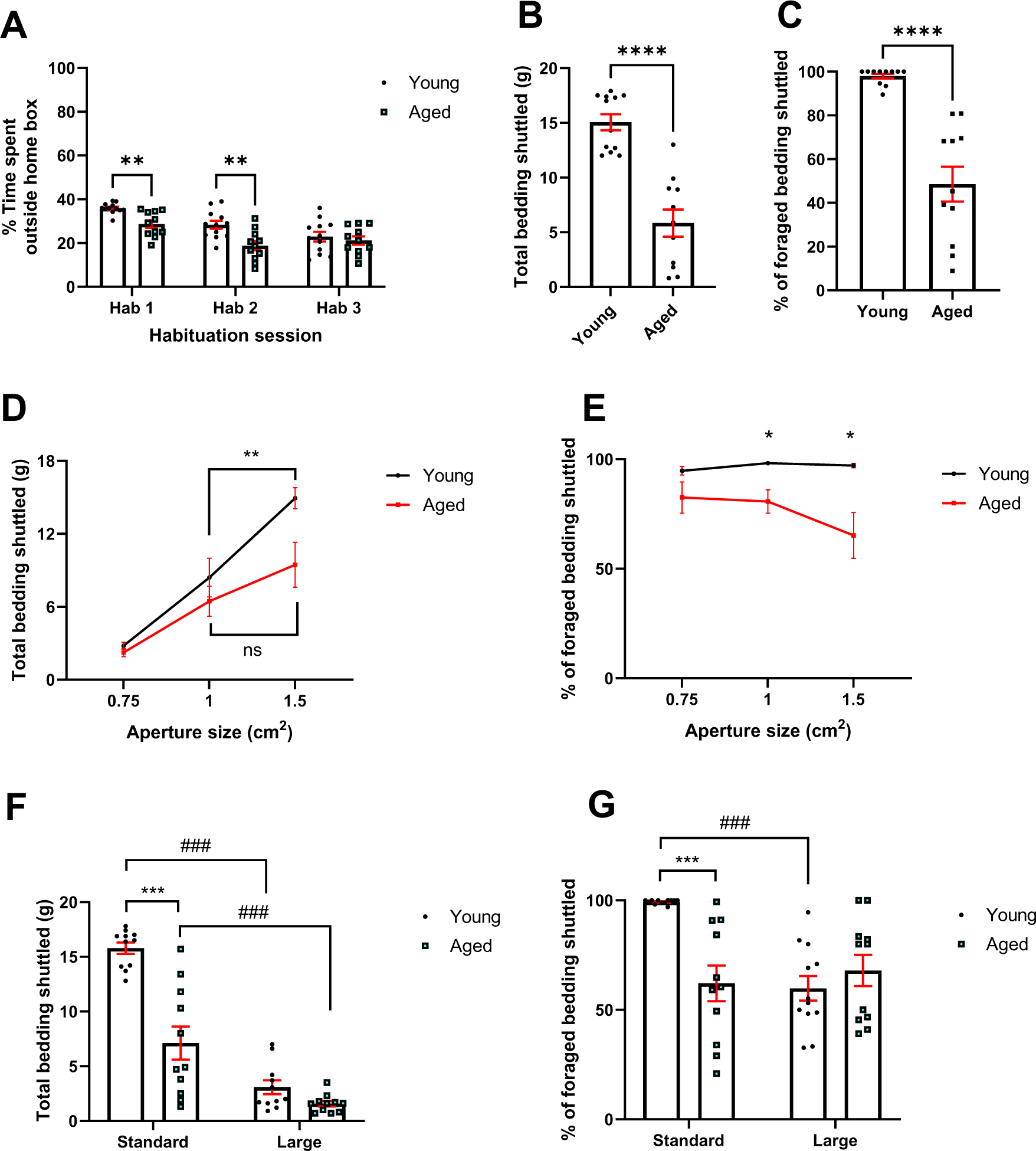
Aged mice show impairments in modulation of foraging behaviour under different effort and affective contingencies. Aged and young mice underwent a series of experiments where aspects of the internal environment were modified. **A** Aged mice showed a reduction in forage area exploration (p < 0.01) until the final session (p > 0.05). **B** In an initial foraging session, aged mice showed both a reduction in total bedding shuttled (p < 0.0001) and **C** % of foraged bedding taken to the home area (p < 0.0001). **D** Reducing aperture size from 1.5 cm^2^ to 1 cm^2^ reduced total bedding shuttled in young but not aged mice (p < 0.01 and p > 0.05 respectively). **E** At both the 1.5 cm^2^ and 1 cm^2^ aperture aged mice showed a reduction in % of foraged bedding taken to the home area (p < 0.05). **F** When the forage area was made larger, both age groups showed a reduction in total bedding shuttled (p < 0.001). Using the standard area, aged mice foraged less than younger mice (p < 0.001) however this effect disappeared in the larger area (p > 0.05). **G** When the forage area was made larger, younger mice left a greater amount of bedding in the forage area (p < 0.001) but aged mice did not (p > 0.05). In the standard area aged mice left more bedding in the forage area than younger mice (p < 0.001) but this effect disappeared in the larger area (p > 0.05). Bars are mean ± SEM, **p<0.01, ***p<0.001, ###p<0.001 (within-subject comparison).

In an initial foraging session, aged mice shuttled less bedding than younger mice (t(21) = 6.505, p < 0.0001) and showed a reduction in % of foraged bedding taken to the home area (t(20) = 6.141, p < 0.0001) (**fig. 8B&C**). Mice then underwent a foraging session under different effort contingencies governed by the size of the bedding box aperture, allowing the formation of an ‘effort curve’. Here, there was a main effect of age (F(1,20) = 4.790, p = 0.0407), a main effect of aperture (F(1.969,39.39) = 47.3, p < 0.0001, and an age*aperture interaction (F(2,40) = 3.242, p = 0.0496) on total bedding shuttled. Sidak’s post hoc analysis revealed that there was a trend towards aged mice foraging less bedding at the widest aperture only (p = 0.0518). Young mice foraged less bedding from the 1 cm^2^ aperture than the 1.5 cm^2^ (p = 0.003) but aged mice did not show this difference (p > 0.05) (**fig. 8D**). In addition, there was a main effect of age on % of foraged bedding taken to the home area (F(1,20) = 9.677, p = 0.0055). There was an aperture*age interaction (F(2,40) = 3.285, p = 0.0478), but no main effect of aperture alone (p > 0.05). Post hoc analysis revealed that aged mice had a lower % of bedding taken back to the home area compared to younger mice at both the 1 cm^2^ aperture and the 1.5 cm^2^ aperture (p = 0.0247 and p = 0.0361 respectively). There was no difference between aperture sizes in either age group (**fig. 8E**).

Next, the impact of the size of the foraging area on foraging behaviour was assessed, based on the principle that rodents find more open spaces more aversive (Rex et al., 1998). Analysis of total bedding taken through revealed a main effect of forage area size (F(1,20) = 104.8, p < 0.0001), a main effect of age (F(1,20) = 36.40, p < 0.0001) and an age*size interaction (F(1,20) = 16.15, p = 0.0007). Post hoc analysis showed that in the standard forage area, aged mice shuttled less bedding than younger mice (p < 0.0001) however this difference disappeared when the forage area was made larger (p > 0.05). Both young and aged mice foraged less in the larger versus standard forage area (p < 0.0001 and p = 0.0006 respectively) (**fig. 8F**). When considering the % of foraged bedding taken to the home area, there was a main effect of age (F(1,21) = 4.416, p = 0.0478, a main effect of forage area size (F(1,21) = 12.20, p = 0.0022) and a size*age interaction (F(1,21) = 22.19, p = 0.0001). Post hoc analysis revealed that in the standard forage area, aged mice showed a lower % of bedding taken back to the home area (p = 0.0001) but not in the larger area. Younger mice had a lower % of foraged bedding taken back to the home area in the larger area versus the standard area, but aged mice did not (p < 0.0001 and p > 0.05 respectively) (**fig. 8G**).

The effect of a chronic corticosterone treatment on foraging behaviour was similarly analysed. In an initial study, CORT treated mice shuttled less bedding than non-treated mice (t(29) = 2.853, p = 0.0079) (**fig. 9A**) but there was no difference in % of foraged bedding taken back to the home area (p > 0.05) between treatment groups (**fig. 9B**). When assessed using the ‘effort curve’ there was a main effect of aperture (F(1.352, 37.85) = 72.83, p < 0.0001), a main effect of CORT treatment (F(1,28) = 5.981, p = 0.021) and an aperture*treatment interaction (F(2,56) = 3.274, p = 0.045) on bedding shuttled (**fig. 9C**). Post hoc analysis revealed that at the 1 cm^2^ aperture, CORT treated mice reduced total bedding shuttled (p = 0.0234). In terms of % of foraged bedding taken to the home area, there was a main effect of aperture (F(1.86,53.02) = 3.515, p = 0.040) and an aperture*treatment interaction (F(2,57) = 5.534, p = 0.0064) but no main effect of treatment (p > 0.05). Post hoc analysis showed that there was no difference in % of bedding taken to the home area between groups at any aperture, though there was a trend towards a reduction in the CORT treated group at 1 cm^2^ (p = 0.0638). Aperture size did not affect % of foraged bedding taken to the home area in the non-treated group, but in the treated group there was a reduction at 1 cm^2^ compared to both 0.75 cm^2^ and 1.5 cm^2^ (p = 0.015 and p = 0.0345 respectively).

**Figure 9.**
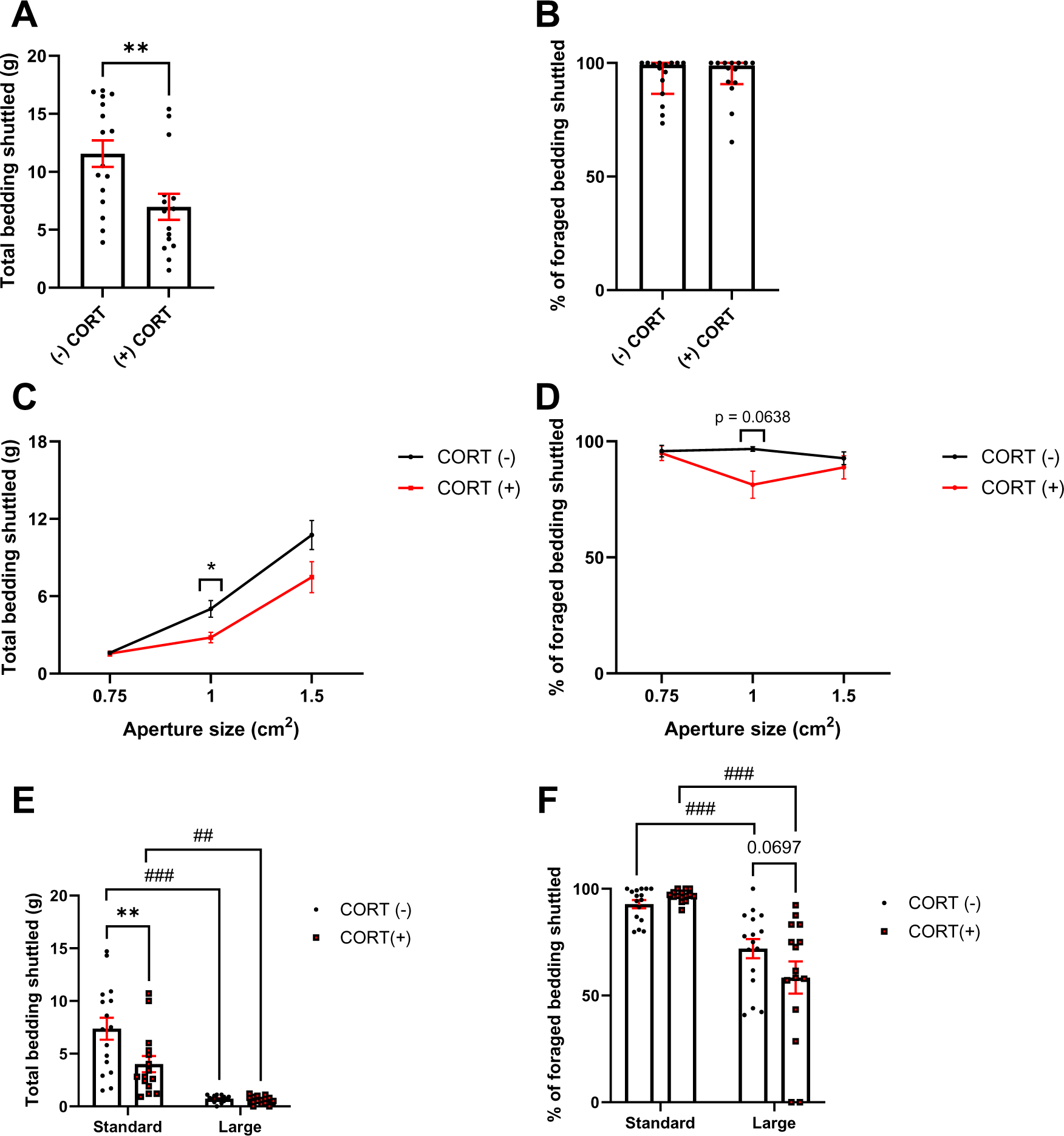
CORT-treated mice show impairments in modulation of foraging behaviour under different effort and affective contingencies. CORT treated and non-treated mice underwent a series of experiments where aspects of the internal environment were modified. **A** In an initial foraging session, CORT treated mice showed a reduction in total bedding shuttled (p < 0.01) **B** but not % of foraged bedding taken to the home area (p > 0.05). **C** Reducing aperture size reduced total bedding shuttled in both groups, and CORT treated mice shuttled less than non-treated mice at the 1 cm^2^ aperture. **D** CORT treated mice did not show any difference in % of foraged bedding taken to the home area at any of the apertures (p > 0.05). **E** When the forage area was made larger, both treated and non-treated groups showed a reduction in total bedding shuttled (p < 0.01 and p < 0.001). Using the standard area, aged mice shuttled less bedding than younger mice (p < 0.01) however this effect disappeared in the larger area (p > 0.05). **F** When the forage area was made larger, both groups left a greater amount of bedding in the forage area (p < 0.01). Bars are mean ± SEM, **p<0.01, ***p<0.001, ##p < 0.01, ###p<0.001 (within-subject comparison).

**Figure 10.**
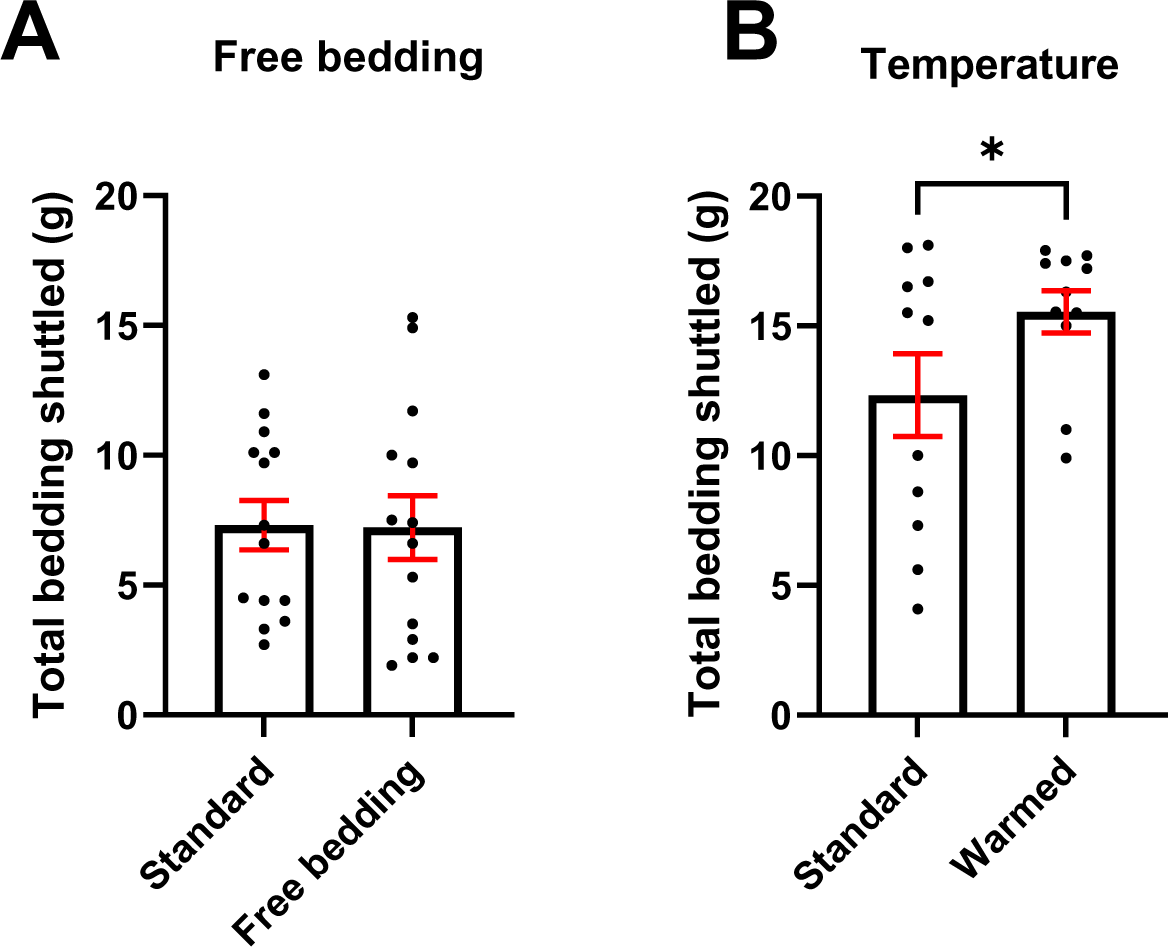
Modulation of aspects of the forage environment change foraging behaviour. Aspects of the forage environment were changed and resulting impact on forage behaviour was analysed. **A** There was no effect of freely available bedding on total bedding shuttled (p > 0.05). **B** Warming the environment increased the amount of bedding shuttled (p < 0.05). Bars are mean ± SEM, * p < 0.05.

Assessing the impact of forage area size on total bedding foraged taken to home area revealed a main effect of aperture (F(1,29) = 63.70, p < 0.0001), a main effect of treatment (F(1,29) = 6.399, p = 0.0171) and an size*treatment interaction (F(1,29) = 6.619, p = 0.0155). Post hoc analysis found that CORT treatment reduced total bedding shuttled in the standard (p = 0.0013) but not large forage area (p > 0.05). Both CORT treated and non-treated groups foraged less in the larger forage area compared to the standard (p < 0.0001 and p = 0.0015 respectively). When % of foraged bedding taken to the home area was compared, there was a main effect of size (F(1,29) = 50.34, p < 0.0001) and an aperture*treatment interaction (F(1,29) = 4.409, p = 0.0446) but no main effect of treatment (p > 0.05). In the large foraging area, there was a trend towards the CORT treated mice showing a reduction in % of foraged bedding taken to the home area compared to non-treated mice (p = 0.0697). In both the treated and non-treated group, a larger area reduced % of foraged bedding taken to the home area (p< 0.0001 and p = 0.0024 respectively).

To investigate the driving factors of foraging behaviour, aspects of the forage environment itself were modulated. Providing 10g of freely available sizzle bedding in the main compartment of the forage arena had no effect on total transported bedding (p > 0.05). Warming the main compartment of the forage area increased the amount of bedding shuttled (t(10) = 2.398, p = 0.0375).

## Discussion

Systemic administration of haloperidol and amphetamine (i.p) decreased bedding shuttled, while methylphenidate increased bedding shuttled. Oral administration of amphetamine had no effect. Healthy ageing reduced bedding shuttled, blunted effort modulation in the effort curve study and a blunted change in shuttle behaviour in a more aversive (more open) environment. Chronic corticosterone similarly reduced bedding shuttled and showed a particular sensitivity to the effort induced by the moderate difficulty aperture (1 cm^2^). Provision of free bedding did not affect bedding shuttled and warming the environment increased foraging behaviour. The following discussion will consider how these findings contribute to our understanding of this novel non-appetitive task as a translational measure of motivated behaviour in mice.

### Foraging behaviour is sensitive to acute dopaminergic modulation

Consistent with appetitive effort-based conditioning paradigms such as the EfR task, foraging behaviour is sensitive to a reduction in DA transmission mediated by the D2 receptor antagonist haloperidol. Acute treatment with 0.1 mg/kg haloperidol reduced the number of high effort, high value reward trials in the EfR task in both mice and rats (Marangoni et al., 2023, Griesius et al., 2020, Salamone et al., 1994). Here, we found a reduction in bedding shuttled to the home box independent of any changes to general locomotion. Similarly, we found a robust increase in bedding shuttled with acute treatment with methylphenidate (MPH) which increases DA signaling via inhibition of the dopamine transporter (DAT). This similarly aligns with previously published EfR results where MPH trended towards an increase in number of high effort, high value reward trials in mice (Marangoni et al., 2023) and has shown similar effects on effort-based decision making in humans (Addicott et al., 2019). Interestingly, treatment with 0.1 mg/kg of the psychostimulant amphetamine (i.p) reduced foraging behaviour which is at odds with findings reported in operant effort-based decision-making tasks where an increase high effort, high reward responding was found (Marangoni et al., 2023, Griesius et al., 2020, Floresco et al., 2008). This divergence in behavioural findings may be driven by differences in task environment. The operant box is a contrived environment designed to encourage robust and trial-restricted behavioural response. Conversely the EBF task environment is more complex, unconstrained by trials and open to a myriad of behavioural options. Low dose amphetamine increases engagement in a variety of different tasks by stabilizing task neural trajectories. This makes it difficult to shift between different behaviours within-task (Hashemnia et al., 2020). These effects are conducive to a contrived environment where behavioural options are limited, but may be impairing in more complex environments. Conversely, oral treatment with amphetamine had no effect on foraging behaviour. This suggests that the initial sharp peak in drug plasma concentration achieved by i.p administration is impairing in this context. It has previously been shown that speed of delivery of DA modulators engage distinct neural circuits (Manza et al., 2023). This suggests the EBF task is sensitive to differing neurotransmitter dynamics mediated by drug pharmacokinetics in addition to absolute changes in DA levels.

By asking animals to engage in behaviours based on intrinsic drive, we have greater access to individual variability, which is often lost under conditions of food restriction or external modulation of motivational state. Stratification of mice into low and high motivation groups revealed that mice in the lower motivation group were more sensitive to lower doses of MPH compared to higher motivated mice. Similarly, stratification revealed that amphetamine (i.p) was impairing only in the higher motivation group. This may be driven by differing basal levels of dopamine within motivation group which provide differing thresholds for dopaminergic drug response. It is important to note that the drugs used do not act upon the dopaminergic system alone, and may also modulate other systems including the noradrenergic system. Use of specific inhibitors are necessary to parse the role of these systems on motivated behaviour more effectively.

Together these results suggest that similarly to effort-based operant conditioning paradigms, the EBF task is sensitive to changes in DA modulation. This provides initial evidence that foraging behaviour may engage aspects of the mesolimbic DA system. These drug effects were achieved without the need to food restrict the mice or required prolonged periods of training. Therefore this task may represent a refinement in the rodent lifetime experience, in addition to accelerating drug screen throughput.

### Foraging behaviour is sensitive to phenotypic models of reward motivation deficit

While the use of acute pharmacology can give us some mechanistic insight and provide evidence for the predictive validity of a model, it cannot fully recapitulate the complexity of a psychiatric disorder. Phenotypic models may therefore provide valuable information as to whether a task can capture relevant behaviours in the context of disease. Normal ageing has been shown to induce behaviours relevant to apathy syndrome including a deficit in reward motivation measured by the EfR task (Jackson et al., 2021, Ishii et al., 2009). Here, we find normal ageing reduced bedding shuttled in an initial foraging session at the widest aperture. In addition, unlike mice that had undergone acute DA modulation, aged mice also showed a reduction in the percentage of bedding foraged shuttled to the home box. This means that of the bedding removed from the bedding box, less was transported to the home box. The significance of leaving foraged bedding in the forage area is currently unclear. However, transportation of bedding may reflect an additional effortful component of foraging behaviour.

Unlike younger mice, aged mice did not adjust their foraging behaviour in response to a more effortful aperture (1.5 cm^2^ vs 1 cm), suggesting a blunting in effort-based behavioural modulation. Consistent with this, previous work has shown that apathetic patients in a Parkinson’s disease patient population showed a reduction in effort sensitivity in a physical effort based decision making task (Le Heron et al., 2018). Of note, percentage of bedding shuttled to the home box was differentially modulated by effort in the aged group. This provides further evidence shuttling of bedding following foraging is additionally effortful. Greater levels of bedding are foraged at the moderate and easy apertures and thus require greater volumes to transport.

In addition to a reduction in reward motivation, apathy syndrome also has an emotional-affective component, where affective reactivity becomes blunted, including response to aversive stimuli (Jackson and Robinson, 2022). Both younger and aged mice showed a reduction in bedding shuttled when the bedding box was placed in a larger (and hence more open) forage area. It is well known that rodents find open environments aversive and are often used as a model of anxiety-like behaviour in various paradigms including the open field test and novelty suppressed feeding test (Rex et al., 1998, Samuels and Hen, 2011). Thus, stress induced by an aversive environment is likely to have suppressed foraging behaviour. Along with a reduction in bedding foraged, younger mice also showed a reduction in percentage of foraged bedding shuttled in the large forage area context, suggesting this measure has an additional affective component. Potentially, transportation of bedding increases time spent in the open which the mouse tries to minimize. Of note, this measure was unaffected in the aged mice irrespective of environment which may be reflective of affective blunting consistent with an apathy phenotype.

Chronic treatment with corticosterone is thought to mimic chronic stress or reflect hypercortisolemia found in patients with major depressive disorder (MDD). It has been shown to induce deficits in reward-related domains and exaggerated emotional reactivity relevant to MDD. However evidence for its effects on reward motivation are mixed. Use of operant based methods of effortful decision making reveal little to no effect (Marangoni et al., 2023, Dieterich et al., 2019). However, an initial foraging session showed a reduction in bedding shuttled in the CORT treated group with no change in percentage of foraged bedding shuttled. Crucially, this suggests that the EBF task is sensitive to behavioural changes that would otherwise go undetected by an operant based approach. Other non-operant methods have also found more robust behavioural effects relating to motivation. Robust effects of CORT treatment in mice were also found in a T-maze task, where a reduction in high effort, but high value reward arm choices was found (Dieterich et al., 2019). This suggests that more ethologically relevant, non-conditioned tasks may be a more sensitive readout of motivational state in some contexts.

Unlike ageing, CORT treatment did not alter behavioural modulation in response to effort requirement, but led to a greater reduction in bedding shuttled when the aperture was moderately challenging. Consistent with above, a more open environment suppressed bedding shuttled in treated and non-treated groups. However, CORT treated mice trended towards a greater reduction in percentage of bedding shuttled suggestive of exaggerated affective reactivity consistent with MDD than apathy syndrome (Jackson and Robinson, 2022). Thus, by changing aspects of the foraging environment we may parse different behavioural profiles relating to motivation and affect from distinct disease phenotypes.

Together, these results suggest that the EBF task can sensitively detect motivational deficit in phenotypic models. It can detect behavioural deficits that may go undetected by the standard operant box approach and can provide distinct behavioural profiles relating to both positive and negatively valanced domains between disease states.

### Foraging behaviour is not sensitive to external changes in bedding value

In contrast to food motivated tasks where food is removed before task onset to induce a robust, conditioned behavioural response, here the mouse is unrestricted and can choose to behave depending on intrinsic motivational state. However, it is important to note that mice construct nests for thermoregulation. It has previously been reported that animal units are often colder than the mouse’s thermoneutral zone of 29 °C (Vialard and Olivier, 2020) and, compounded by the fact that the mice are singly housed both in and outside of the task, may be relatively cold. Thus, the observed foraging behaviour may be driven by a desire to keep warm. If this is the case, warming the environment should devalue the bedding and thus reduce foraging. However, warming the home box to ∼29 °C increased foraging behaviour. This suggests that warming the environment increased activity but did not devalue the bedding ‘reward’. However, to assess this comprehensively a temperature ‘bell curve’ should be generated to assess the impact of a broader range of external temperatures on motivation to perform this behaviour. Similarly, providing a large quantity of free sizzle bedding in the home box where nest construction occurs should also devalue the bedding available in the bedding box which requires additional effort to obtain. However, provision of free bedding did not reduce bedding shuttled. Together these data suggest that it is the act of foraging itself that is rewarding in this context, rather than the retrieval/use of the bedding itself. This contrasts with existing operant paradigms where the rewarding aspect of the task is the consumption of a food reward following the exertion of effort. In this case, foraging behaviour may be an intrinsically motivating, or ‘self-driven behaviour’. This behaviour may therefore be more relevant to apathy, which is characterised by a reduction in self-initiated behaviour. It is less clear whether this type of deficit can be detected under the pressure of a significant external motivator such as hunger.

It is important to acknowledge that this study used males only and greater translatability would be achieved with the use of both sexes. There is the potential for sex-related differences in motivated foraging behaviour which will be assessed in future studies.

## Conclusion

Using acute dopaminergic modulation along with two phenotypic models of motivational deficit we investigated the utility of the EBF task as a measure of motivational state in mice. We show that in line with other effortful decision making paradigms, the EBF is sensitive to DA manipulation suggesting some involvement of the mesolimbic DA system. Using healthy ageing and chronic corticosterone treatment we demonstrate the EBF task is a sensitive measure of motivational deficit and potential affective changes in relevant phenotypic models previously undetected with EfR methods. We found that foraging behaviour was not reduced when the bedding was devalued, suggesting engaging in foraging itself is intrinsically rewarding. Thus, non-appetitive foraging represents a complex behaviour that offers a unique perspective on intrinsic motivational state. Asking animals to exert effort via more ethologically-relevant behaviours may have greater translational value, particularly in the context of apathy drug discovery where self-driven behaviours may play a vital role. While people don’t leave their houses to forage for bedding per se, this task reflects the engagement of an animal with its own environment. The animal is not performing the same action in the same context as a human, but simply engaging in a motivated behaviour relevant to its own ethology.

## Acknowledgments

The authors gratefully acknowledge Julia Bartlett for technical assistance in behavioural testing, Professor Emma Robinson for provision of experimental space and support, and the University of Bristol Animal Services Unit for care of the mice.

## Data accessibility

All data used in this manuscript will be made accessible on the Robinson Lab Open Science Framework upon acceptance.

