## Supplementary material for "Validation of a non-appetitive effort-based foraging task as a measure of motivational state in male mice"

**S1**

| Experiment | Length | Aperture size | Cohort | N number | Age |
| --- | --- | --- | --- | --- | --- |
| Free bedding | 2h | Mid | MJ16 | 16 | 35 weeks |
| Temp study | 4h | Large | MJ15 | 12 | 30 weeks |
| Age: First session | 4h | Large | MJ15/JD5 | 12/group | 13 weeks (Y) 44 weeks (O) |
| Age: Effort curve | 4h | All | MJ15/JD5 | 12/group | 23 weeks (Y) 54 weeks (O) |
| Age: Big vs std | 4h | Large | MJ15/JD5 | 12/group | 19 weeks (Y) 50 weeks (O) |
| CORT: First session | 4h | Large | MT1/BR1 | 16/group | BR1 35 weeks, MT1 18 weeks |
| CORT: Effort curve | 2h | All | MT1/BR1 | 16/group | BR1 35 weeks, MT1 19 weeks |
| CORT: Big vs std | 2h | Large | MT1/BR1 | 16/group | BR1 37 weeks, MT1 21 weeks |
| Haloperidol i.p | 2h | Mid | MJ16 | 16 | 12 weeks |
| Amphetamine i.p | 2h | Mid | MJ16 | 16 | 14 weeks |
| Amphetamine oral | 2h | Mid | MJ17 | 16 | 15 weeks |
| Methylphenidate oral | 2h | Mid | MJ17 | 16 | 13 weeks |

***S1. Summary of n numbers and ages per experiment.***

**S2**

| **Component** | **Dimensions and links** |
| --- | --- |
| Home area | W 17.5 cm, L 30.0 cm, H 13.0 cm |
| Home area lid | W 18.1 cm, L 18.1 cm, H 18.1 cm |
| Connecting tube | L 21.0 cm |
| Forage area | W 10.0 cm, L 10.0 cm, H 13.0 cm |
| Forage area (large) | W 21.0 cm, L 21.0 cm, H 14.0 cm |
| Bedding box body (3D printed) | W 3.5 cm, L 8.0 cm, H 10.0 cm |
| Bedding box face plate (3D printed) | W 0.3 cm, L 7.8 cm, H 10.0 cm |
| Bedding box lid (3D printed) | W 3.5 cm, L 7.8 cm, H 0.3 cm |
| Magnet bar (3D printed) | W 2.0 cm, L 8.0cm, H 3.0 cm |

***S2. Component dimensions and 3D print files for forage arena.***

**S3**

| Experiment | Measure | Exclusions in dataset |
| --- | --- | --- |
| Free bedding | Total taken through | N = 1, std, outlier |
| Temp study | Total taken through | N = 1 pre-exclusion |
| Age: Habituation | Time spent in box(s) | N = 2 young, hab 1 (n = 1 missing value), n = 1 aged, hab 3 |
| Age: First session | Total taken through | No exclusions |
|  | % taken to main box | N = 1 young |
| Age: Effort curve | Total taken through | N = 1 pre-exclusion, n = 1 young, 15cm outlier |
|  | % taken to main box | N = 1 pre-exclusion, n = 1 young 0.75cm, n = 1 aged 1cm, outliers |
| Age: Big vs std | Total taken through | N = 1 pre-exclusion, n = 1 big, young, n = 1 big, aged, outliers |
|  | % taken to main box | N = 1 pre-exclusion, n = 1 std, young |
| CORT: First session | Total taken through | No exclusions |
|  | % taken through | N =1 (-) n = 1 (+) outliers |
| CORT: Effort curve | Total taken through | N = 1 pre-exclusion due to box error.  N = 1 0.7cm (-), n = 1 1cm (-), n = 1 1.5cm (+). |
|  | % taken to main box | N = 1 foraged 0g so % could not be calculated. |
| CORT: Big vs std | Total taken through | N = 1 big (-), n = 1 big (+), n = 1 std (+) outliers |
|  | % taken to main box | N = 1 big (-), n = 1 std (+) outliers |
| Haloperidol | Total taken through | N = 1 vehicle (box error) |
|  | Performance split | N = 1 0.03mg/kg outlier |
|  | % taken to main box | N = 2 0.01mg/kg, n = 2 0.03 mg/kg, n = 1 0.1 mg/kg outliers. |
|  | Activity | N = 3 0.03mg/kg sensor error |
| Amphetamine i.p | Total taken through | N = 1 0.1 mg/kg outlier  N = 2 0.3 mg/kg outlier, missing value. |
|  | Performance split | N = 1 0.1 mg/kg. |
|  | % taken to main box | N = 1 excluded due to 3 outliers.  N = 1 0.1 mg/kg, n = 1 1.0mg/kg outliers. |
|  | Activity | N = 1 vehicle, n = 1 0.01 mg/kg, n= 2 0.3 mg/kg, n = 1 1.0mg/kg outliers |
| Amphetamine oral | Total taken through | N = 1 vehicle, outlier |
|  | % taken to main box | N = 1 vehicle, n = 1 0.1mg/kg, n = 1 0.3mg/kg outliers |
|  | Split | No exclusions |
| MPH oral | Total taken through | N = 2, 10 mg/kg, n = 1 vehicle, outliers |
|  | % taken to main box | N = 1 fully excluded due to multiple outlying points, n = 1 10mg/kg outlier. |
|  | Split | N = 1 1mg, high, outlier |

***S3. Summary of data point exclusions/replacements per experiment.***
